# Foxn1 overexpression promotes thymic epithelial progenitor cell proliferation and mTEC maintenance, but does not prevent thymic involution

**DOI:** 10.1101/2022.06.01.494345

**Authors:** Jie Li, Lucas P. Wachsmuth, Shiyun Xiao, Brian G. Condie, Nancy R. Manley

## Abstract

The transcription factor FOXN1 is essential for fetal thymic epithelial cell (TEC) differentiation and proliferation. In the postnatal thymus, *Foxn1* levels vary widely between different TEC subsets, from low or undetectable in putative TEC progenitors to highest in the most differentiated TEC subsets. Correct *Foxn1* expression is also required to maintain the postnatal microenvironment, as premature down-regulation of *Foxn1* causes a rapid involution-like phenotype, while transgenic over-expression can cause thymic hyperplasia and/or delayed involution. In the current study, we investigated a *K5.Foxn1* transgene that drives *Foxn1* over-expression in TECs, but does not cause hyperplasia, nor does it delay or prevent aging-related involution. Similarly, this transgene cannot rescue thymus size in *Foxn1^lacZ/lacZ^* mice that undergo premature involution due to reduced *Foxn1* levels. However, *K5.Foxn1* transgenics do maintain TEC differentiation and cortico-medullary organization with aging both alone and in hypomorphic *Foxn1^lacZ/lacZ^* mice. Analysis of candidate TEC markers showed co-expression of progenitor and differentiation markers as well as increased proliferation in Plet-1+ TECs associated with *Foxn1* expression. These results demonstrate that the functions of FOXN1 in promoting TEC proliferation and differentiation are separable and context-dependent, and suggest that modulating *Foxn1* levels can regulate the balance of proliferation and differentiation in TEC progenitors.

## Introduction

The thymus provides the essential microenvironment for lineage commitment and development of T cells. Thymic epithelial cells (TECs) include medullary (mTEC) and cortical (cTEC) cells that mediate the homing of lymphoid progenitors and the proliferation, survival, and differentiation of developing T cells. TECs derive from the 3rd pharyngeal pouch endoderm during fetal development, which proliferates to form the thymic rudiment at about E11.5 in mouse embryonic development (Gordon and Manley, 2011). In both humans and mice, the thymic rudiment becomes functional only after the transcriptional activation of the *Foxn1* gene in the thymic epithelium (Brissette et al., 1996; Romano et al., 2013).

FOXN1 is a key transcription factor that controls most aspects of TEC proliferation and differentiation. *Foxn1* is selectively expressed in thymic and skin epithelia, where it regulates the expression of downstream molecular targets to control both growth and differentiation (Brissette et al., 1996). In mice, rats (Nehls et al., 1994), and humans (Romano et al., 2012) null mutations in the *Foxn1* gene (nude) lead to a hairless phenotype and alymphoid cystic thymic dysgenesis due to defective TEC differentiation (Brissette et al., 1996; Nehls et al., 1996; Nehls et al., 1994). In the skin, FOXN1 promotes keratinocyte proliferation and suppresses differentiation, and must be down-regulated for differentiation to proceed (Li et al., 2007). In contrast, in fetal and postnatal TECs FOXN1 is required both for proliferation and for progression of differentiation at multiple stages in cTEC and mTEC sub-lineage development (Nowell et al., 2011; Su et al., 2003; Vaidya et al., 2016). TECs are exquisitely sensitive to FOXN1 levels, and even small changes in FOXN1 dose can affect phenotypes (Chen et al., 2009; Nowell et al., 2011). Furthermore, maintenance of FOXN1 levels is required for postnatal thymus homeostasis. In *Foxn1^lacZ^* mutant mice, premature postnatal down-regulation of *Foxn1* causes disorganization and atrophy of the thymus similar to premature aging-related thymic involution (Chen et al., 2009) while two different models of transgenic over-expression of *Foxn1* exhibit prolonged maintenance of thymic size, structure, and function, delaying involution (Bredenkamp et al., 2014a; Zook et al., 2011).

It is as yet unclear what the role of FOXN1 is in TEC progenitor/stem cells (TEPCs/TESCs). This uncertainty is in part due to lack of clarity on the unique phenotypes and functional capacity of both fetal and postnatal TEPC. In the fetal thymus, the EpCam+Plet1+ TEC population includes a common thymic epithelial precursor (TEPC), from which both cTECs and mTECs will be subsequently generated (Bennett et al., 2002; Gray et al., 2006). As these cells were originally identified based on their phenotype in *Foxn1* null nude mice (Blackburn et al., 1996), it is possible that TESCs are *Foxn1* negative, at least initially. Furthermore, TEPC/TESC can persist for long periods in the absence of or under very low Foxn1 levels, and can be activated by increasing Foxn1 expression (Bleul et al., 2006; Jin et al., 2014). However, there is also evidence that all TECs express *Foxn1* at some point early in their differentiation (Corbeaux et al., 2010; O’Neill et al., 2016). Furthermore, it is clear that during differentiation both cTEC and mTEC lineages modulate *Foxn1* levels over a wide range (Chen et al., 2009; Nowell et al., 2011). FOXN1 may have varying effects at different levels in these distinct populations; for example, there is evidence that lower FOXN1 levels preferentially promote proliferation in less mature MHC Class II low (MHCII^lo^) TECs, whereas higher levels promote differentiation into MHCII^hi^ populations (Bredenkamp et al., 2014a; Chen et al., 2009; Nowell et al., 2011). Taken together, these and other data support a model in which TESCs/TEPCs express at most very low *Foxn1* levels, and that its differential up-regulation controls both TEC proliferation and differentiation at multiple stages of both mTEC and cTEC differentiation. Although understanding these dynamics is critical, neither how *Foxn1* levels are regulated in these different sub-populations nor FOXN1’s specific roles across TEC subsets are yet well understood.

In this study, we investigated the quantitative and TEC subset-specific roles of FOXN1 using *K5.Foxn1* transgenic (Weiner et al., 2007), *Foxn1^lacZ^* (Chen et al., 2009), and nude *(Foxn1* null; (Nehls et al., 1994)) mouse strains. We show that the *K5.Foxn1* transgene drives *Foxn1* expression broadly in TECs, resulting in FOXN1 levels 4-6 fold higher than normal. However, unlike previously published overexpression models, this transgene did not result in increased thymus size or delay thymus size reduction with involution in either wild-type or *Foxn1^lacZ/lacZ^* mice. However, *K5.Foxn1* did improve TEC differentiation and thymus function. Importantly, this transgene drove significantly enhanced *Foxn1* expression in Plet1+ cells that contain putative TEC progenitors. This higher expression resulted in both an expanded Plet1+ population and their increased proliferation, as well as ectopic expression of the mTEC progenitor marker Cld3,4 (Hamazaki et al., 2007) and mTEC differentiation marker UEA-1 within Plet1+ cells. In the absence of endogenous *Foxn1,* the *K5.Foxn1* transgene was sufficient to drive formation of a small thymus, with a bias toward mTEC development and an expanded Cld3,4+ population.

These results show that moderate up-regulation of *Foxn1* in TECs broadly biases them toward differentiation without increasing proliferation. Furthermore, *Foxn1* up-regulation within a TESC/TEPC-containing population causes both increased proliferation and misexpression of mTEC markers. Thus, the effects of increasing *Foxn1* levels are context-dependent, a finding that has important implications for efforts to delay thymic involution or rejuvenate the involuted thymus through manipulation of *Foxn1* levels.

## Results

### Increased *Foxn1* expression in *K5.Foxn1* transgenic mice does not prevent thymic involution

The *K5.Foxn1* transgenic mouse line was developed in the Brissette lab to investigate the role of *Foxn1* in skin and hair development by driving its overexpression using a keratin 5 (K5) promoter (Weiner et al., 2007). As K5 is also expressed in multiple populations of TECs, we evaluated thymus phenotypes across aging. We measured the relative increase in *Foxn1* expression at the mRNA level in mTECs (defined as CD45-EpCam+MHC+UEA1+ cells) and cTECs (defined as CD45-EpCam+MHC+UEA1-cells) from *K5.Foxn1*Tg and wild type mice thymi. We used primers specific for the transgene mRNA product, as well as common primers that amplify both the transgenic and endogenous *Foxn1* transcripts. Total expression of *Foxn1* mRNA was increased in TECs from *K5.Foxn1*Tg mice compared to wild type controls (Fig. 1A). Although expression of the endogenous K5 gene in the thymus is primarily restricted to mTECs, the *K5.Foxn1* transgene drives expression in both cTECs (UEA-1-) and mTECs (UEA-1+). *Foxn1* expression in UEA-1+ mTECs increased more than 6-fold, while expression in cTECs increased approximately 4-fold (Fig. 1B). Overall FOXN1 protein levels as detected by immunofluorescence were also clearly increased in both cortex and medulla in transgenic mice (Fig. 1C). Analysis of fluorescence intensity showed increases in both average fluorescence intensity (Fig. 1D) and the proportion of cells with high FOXN1 levels (Fig. 1E) compared to wild type.

**Fig. 1.**
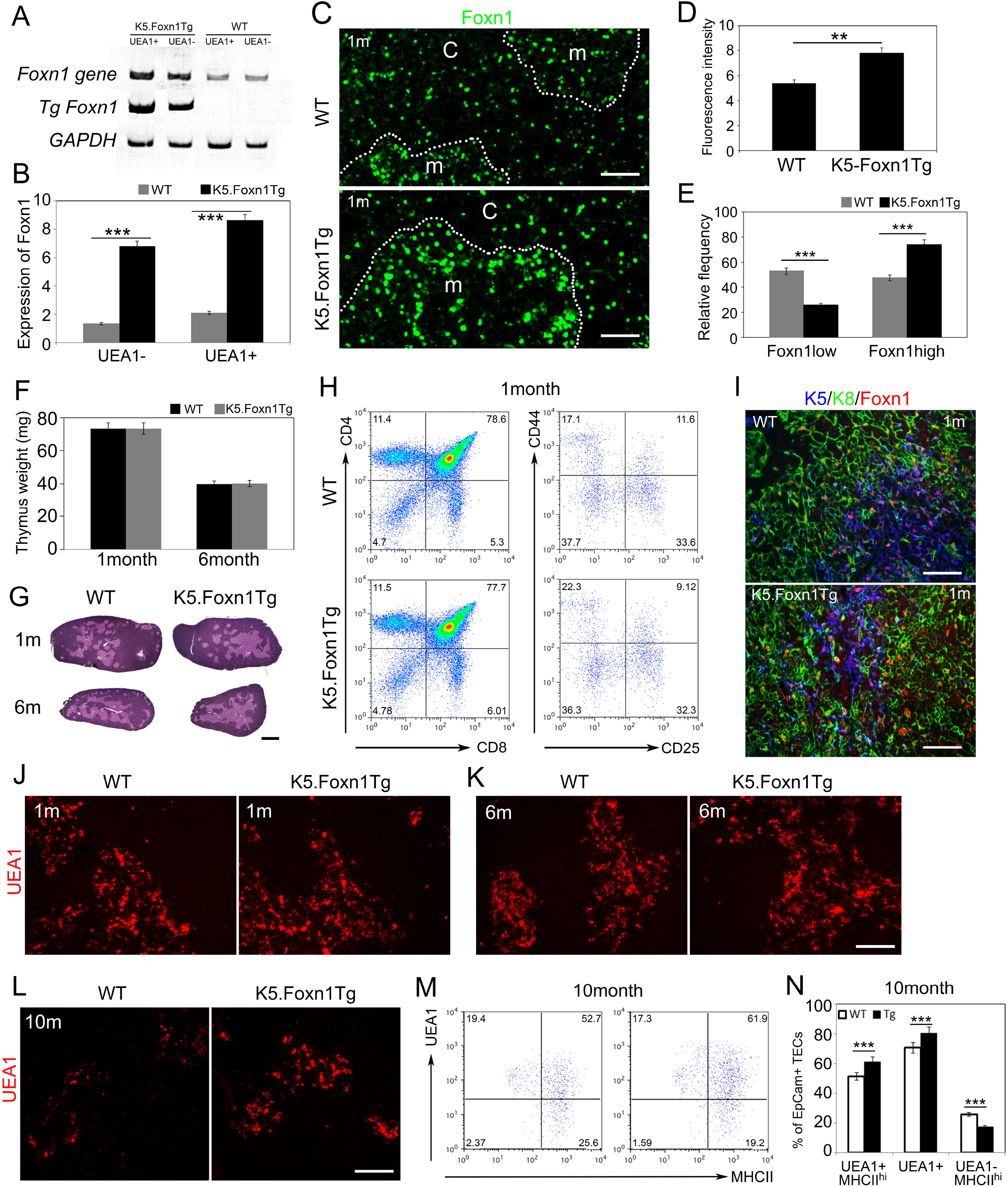
Expression and phenotypes of *K5.Foxn1* in a wild-type background. The *K5.Foxn1* transgene is indicated in all figures as “Tg+”. All ages are 1 month unless otherwise indicated on the figure panels. **(A)** RT-PCR of endogenous- and tg-expressed *Foxn1* in thymic epithelial cells. **(B)** RT-PCR analysis of total *Foxn1* expression in sorted TEC subsets from WT and *K5.Foxn1* thymi. **(C)** Paraffin sections from WT and *K5.Foxn1* Tg+ stained with a FOXN1 antibody (green). **(D)** Quantification of fluorescence intensity of FOXN1 in *K5.Foxn1* transgenic thymus. **(E)** Relative frequency of FOXN1hi cells increased in *K5.Foxn1* TECs. **(F)** Thymus wet weight in *K5.Foxn1* Tg+ and WT mice. **(G)** Hematoxylin- and eosin-stained paraffin sections of thymi from *K5.Foxn1*Tg+ and WT mice. **(H)** FACS profiles of CD4, CD8, CD25, and CD44 expression in *K5.Foxn1*Tg+ and WT thymocytes. **(I)** IHC of K8 (green), K5 (blue), and FOXN1 (red) on *K5.Foxn1*Tg+ and WT thymi. **(J-L)** Fluorescent immunostaining of UEA1 on one **(J),** six **(K),** and 10 month **(L)** *K5.Foxn1*Tg+ and WT thymi. **(M,N)** Flow cytometric analysis of UEA1 shows increased density and percentage of UEA1+ TECs in *K5.Foxn1*Tg mouse thymus compared with WT. Scale bars=100um. n≥5

This level of overexpression was considerably more modest than those observed in previously reported *Foxn1* overexpression models (>20-fold in (Bredenkamp et al., 2014b; Zook et al., 2011)), both of which caused increased thymus size and delayed involution. To investigate the effect of this more moderate level of *Foxn1* overexpression effects on the thymus, we first measured thymus size and weight at 1 month and 6 months. To our surprise, thymus size was not changed by the presence of the transgene (Fig. 1F, G). The overall architecture of the transgenic thymus, as examined by either H&E at 1 and 6 months (Fig. 1G), or by Keratin 5 (K5) and Keratin 8 (K8) staining at 1 month, Fig. 1I) was also not obviously affected. Thymocytes from 1 or 6 month old transgenic mice were similar to wild-type controls in the relative frequencies of CD4+ or CD8+ single-positive (SP), CD4+8+ double positive (DP), CD4-8-double negative (DN) thymocytes, or CD25 and CD44 labeled DN subsets (Fig. 1H, Supplementary Fig. 1).

These data indicated that the modest level of overexpression in this model was insufficient to affect steady-state phenotypes, or to prevent or delay involution in general. However, analysis of UEA-1 staining in 1, 6, and 10 month-old mice showed an increase in the frequency and intensity of UEA-1+ mTEC, both by immunofluorescence (IF) (Fig. 1J-L) and by flow cytometry (Fig. M), with a selective increase in the frequency of UEA-1+MHCII^hi^ cells (Fig. 1N). These data suggested that while neither initial thymus development, steady-state phenotypes, nor involution as measured by thymus size were affected, mTEC differentiation was maintained better with aging compared to wild-type mice.

### K5-Foxn1 transgene partially rescues TEC differentiation phenotypes in *Foxn1^Z/Z^* mice

We have previously reported an allele of *Foxn1, Foxn1^lacZ^*, in which insertion of the *lacZ* gene into the 3’UTR results in postnatal premature down regulation of *Foxn1* to levels about 35% of wild-type (Chen et al., 2009) (referred to hereafter as *Foxn1^Z^*). This early down regulation results in premature thymic involution that is phenotypically similar to but much more rapid than normal aging-related involution, including loss of mTEC markers, reduced MHC Class II (MHCII) expression, and disorganization of cortical-medullary structure (Chen et al., 2009). We tested whether increasing *Foxn1* levels using the *K5.Foxn1* transgene in *Foxn1^Z/Z^* mice would prevent or modulate premature involution in this model. As in the *K5.Foxn1* transgenics on a *Foxn1* wild type background, thymus size was not changed by the presence of the transgene at either one or six months of age in either *Foxn1^+/Z^* (+/*Z*) or *Foxn1^Z/Z^* (*Z/Z*) mice (Fig. 2A, B). Both thymocyte and TEC cellularity at 4 weeks of age were similar regardless of presence of the transgene (Supplemental Fig. 3A, 4A).

**Fig. 2.**
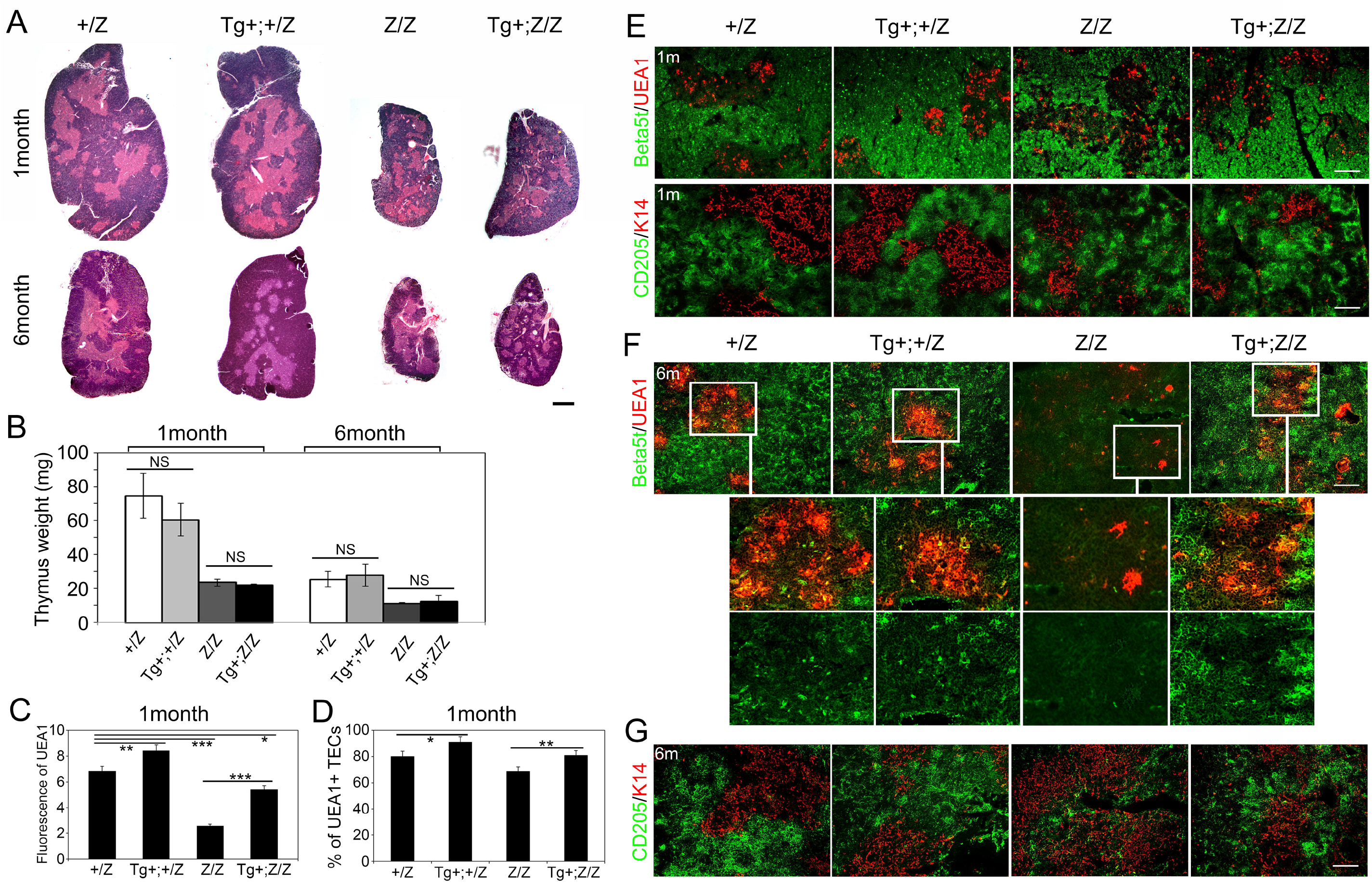
*K5.Foxn1* transgene improves *Foxn1^Z/Z^* induced thymic involution. The *Foxn1^Z^* allele is indicated in all figures as “Z”. All ages are indicated on figure panels. **(A)** Hematoxylin-and eosin-stained paraffin sections of thymi from +/Z, *K5.Foxn1*Tg+;+/Z, Z/Z and *K5.Foxn1*Tg+;Z/Z mice. Scale bar=200μm. **(B)** Thymus wet weight in +/Z, *K5.Foxn1*Tg+;+/Z, Z/Z and *K5.Foxn1*Tg+;Z/Z mice. The *K5.Foxn1* transgene did not induce significant increase in thymus weight. **(C)** Fluorescence intensity of UEA1 in +/Z, *K5.Foxn1*Tg+;+/Z, Z/Z and *K5.Foxn1*Tg+;Z/Z thymus. **(D)** Percentage of UEA1 expressed cells in +/Z, Tg+;+/Z, Z/Z, and Tg+;Z/Z thymic epithelial cells. **(E)** Cryosections from one month old thymi stained for Beta5t (green) and UEA-1 (red) or, CD205 (green) and K14 (red). Scale bar=100μm. **(F)** Cryosections from six month old thymi stained for Beta5t (green) and UEA-1 (red) or, **(G)** CD205 (green) and K14 (red). Scale bars=100μm. n≥5

To further investigate how TEC differentiation in the *Z/Z* mutants was affected by the *K5.Foxn1* transgene, we examined the expression of region-specific markers at 1 and 6 months of age by IHC. For cTECs, we used a cTEC-specific catalytic subunit of thymoproteasome-b5-thymus (β5t) that is a downstream target of FOXN1 (Uddin et al., 2017; Zuklys et al., 2016) and CD205, an early marker of cTEC differentiation (Baik et al., 2013; Jiang et al., 1995). We used both Keratin 14 (K14) and UEA-1 to evaluate mTEC differentiation, which mark largely nonoverlapping mTEC subsets by IHC (Klug et al., 1998).

CD205 was relatively unaffected by changes in *Foxn1* levels in these models at either age (Fig. 2E, G). At 1 month cortical staining for β5t protein was similar between +/*Z* and *Z/Z* thymus in the presence and absence of the *K5.Foxn1* transgene, suggesting that the levels of FOXN1 protein in all of these genotypes was still sufficient for β5t expression (Fig. 2E). However, by 6 months of age β5t was present at very low levels in *Z/Z* thymus compared +/Z and was up regulated in *K5.Foxn1;Z/Z* thymi to levels similar to controls (Fig. 2F).

We previously showed that mTECs in general and UEA-1+ mTECs in particular are decreased in *Z/Z* mutants (Chen et al., 2009). We confirmed this result by both IHC and flow cytometry, in which the fluorescence intensity and % of UEA-1+ mTEC were lower in Z/Z compared to +/Z mice at 1 month of age (Fig. 2C, D; Supplementary Fig 2). Addition of the *K5.Foxn1* transgene increased UEA-1+ levels, in both +/*Z* and *Z/Z* thymi even at 1 month (Fig. 2C). The frequency of UEA1+ cells in both +/*Z* and *Z/Z* thymi were also increased at one 1 month with the addition of the transgene, and frequency in *K5.Foxn1;Z/Z* was rescued to a level similar to +/*Z* mice (Fig. 2D).

Given the early phenotypes in mTECs, we further investigated mTEC differentiation at 1 month by measuring MHCII and AIRE levels. In both +/Z and Z/Z mice presence of the transgene did not affect the frequency of UEA-1+MHCII^hi^ mTECs (Fig. 3A, B). There were however shifts in MHCII^lo^ cells specifically in *K5.Foxn1;Z/Z* mice, with more MHCII^lo^ mTECs and fewer MIIC11^1^” cTECs compared to *Z/Z* (Fig. 3B). As the total number of TECs was not changed in these mice (Supplementary Fig. 3A), this shift in the frequency of MHCII^lo^ mTECs reflected an overall increase in mTEC:cTEC ratio (Fig. 3C). We then evaluated AIRE, a critical mTEC maturation marker, by IHC. AIRE levels decreased more than 5-fold in *Z/Z* mutants on a per cell basis compared to +/Z in one month old mice (Fig. 3D, E), and the frequency declined more than 2-fold (Supplementary Fig. 3B, C). Presence of the transgene did not affect AIRE levels in +/Z mTECs, but in *K5.Foxn1;Z/Z* mice both AIRE levels (Fig. 3D, E) and frequency (Supplementary Fig. 3B,C) were rescued similar to +/*Z* mice. These data are consistent with other reports that *Aire* expression declines with either aging or with declines in *Foxn1* expression (Coder et al., 2015; Xia et al., 2012).

**Fig. 3.**
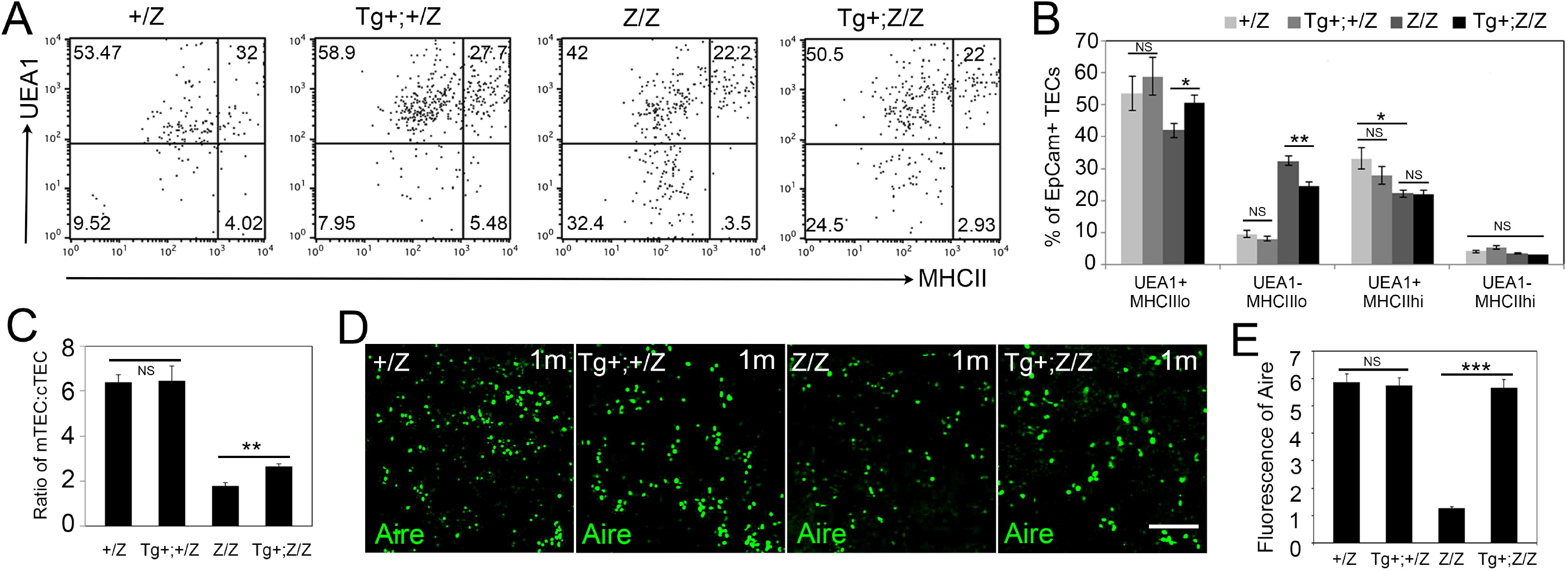
TEC phenotypes in *K5.Foxn1;Foxn1^Z/Z^* mice. All data collected from one month old mice. **(A)** FACS profile of gated CD45-EpCam+ epithelial cells from one month thymi stained for MHCII and UEA-1. **(B)** Frequencies of TEC subsets based on UEA-1 and MHCII staining of EpCam+ TEC from +/*Z*, *K5.Foxn1*Tg+;+/Z, Z/Z and *K5.Foxn1*Tg+;Z/Z thymus. **(C)** Ratio of mTECs:cTECs in +/Z, *K5.Foxn1*Tg+;+/Z, Z/Z and *K5.Foxn1*Tg+;Z/Z thymus. **(D)** Cryosections stained for AIRE (green). The number and intensity of AIRE+ cells were decreased in Z/Z mutants, and rescued by transgene expression. Scale bar=50μm. **(E)** Fluorescence intensity show AIRE levels were rescued by the *K5.Foxn1* transgene in Z/Z thymus. n≥5

These data suggest that maintaining and/or increasing *Foxn1* expression using the *K5.Foxn1* transgene prevented the declines in both cTEC and mTEC differentiation in *Foxn1^Z/Z^* mice, with a more substantial positive impact on mTECs.

### Thymocyte development in *Foxn1^Z/Z^* mice is partially rescued by the *K5.Foxn1* transgene

In *Foxn1^Z/Z^* mutant mice, the TEC defect results in rapid, cell non–autonomous defects in thymocyte development, including reduced total cellularity, selective reduction in CD4+ single positive (SP) cells, and decreased DN1a,b/ETP cells (Chen et al., 2009). Since presence of the transgene did impact TEC differentiation, we tested whether the presence of the transgene in +/*Z* and *Z/Z* mice improved thymocyte differentiation at 1 and 6 months. Similar to wild-type mice (Fig. 1H), in +/*Z* heterozygotes presence of the transgene did not significantly change total thymocyte numbers or the numbers or relative frequencies of CD4-8-double negative (DN), CD4+8+ double positive (DP), or CD4+ or CD8+ SP thymocytes (Fig. 4A, B; Supplemental Fig. 4A-C, E, F), or of the DN subsets defined by CD44 and CD25 (DN1-4) (Fig. 4C, D; Supplemental Fig. 4G, H). In *K5.Foxn1;Z/Z* mice, the numbers and frequencies of the DN and DP subsets were also unaffected, although those of both CD4 and CD8 SP thymocytes were increased slightly, but significantly, relative to *Z/Z* alone at both ages (Fig. 4A, B; Supplemental Fig. 4C).

**Fig. 4.**
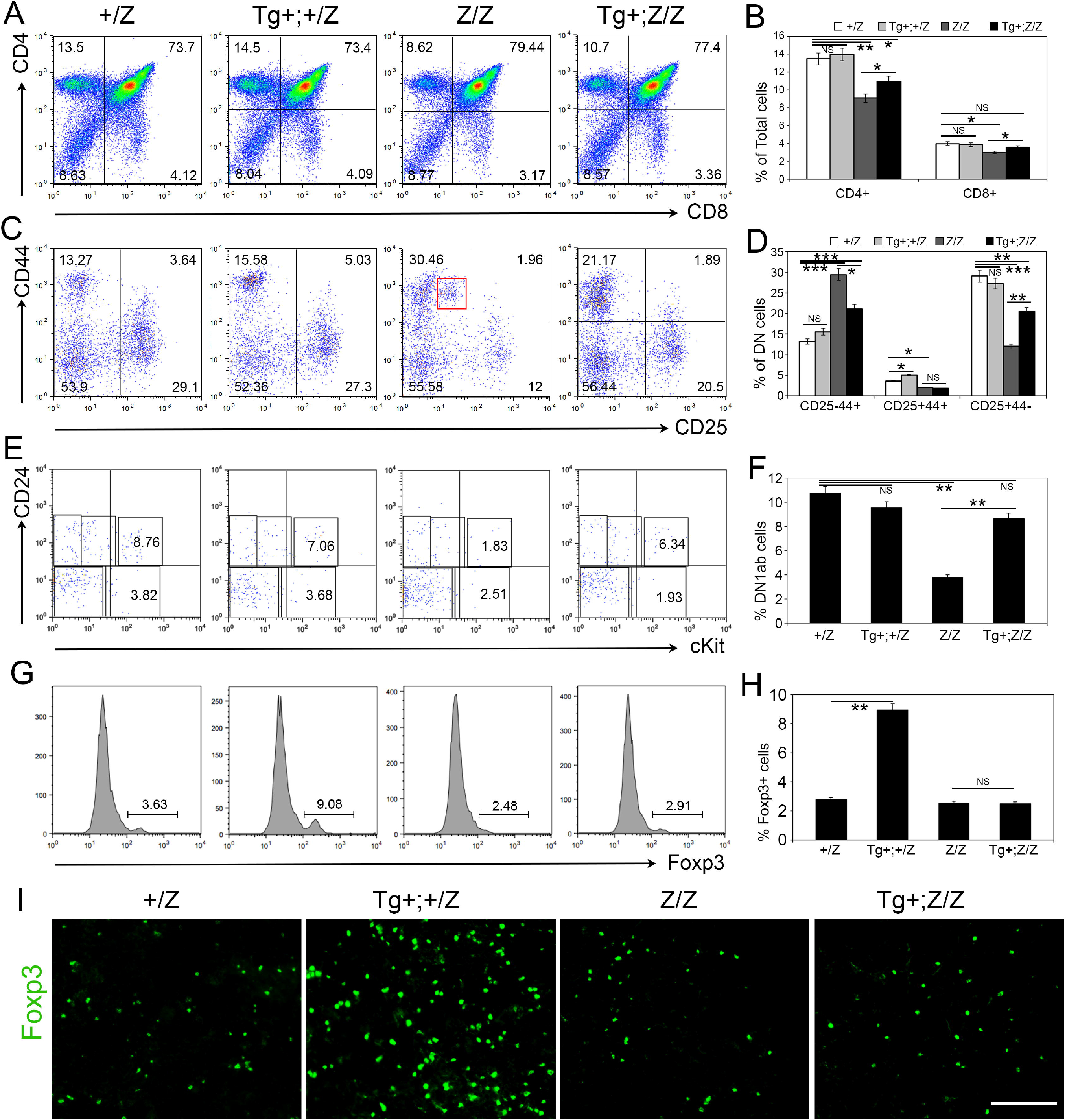
Thymocyte phenotypes were partially rescued by *K5.Foxn1* expression in *Foxn1^Z/Z^* mutants. All mice are 6 months old. **(A)** Profile of CD4 and CD8 expression in +/Z, Tg+;+/Z, Z/Z and Tg+;Z/Z thymocytes. **(B)** The percentage of CD4 and CD8 expressed cells in +/Z, Tg+;+/Z, Z/Z and Tg+;Z/Z thymocytes. **(C)** Profile of CD25 and CD44 expression in gated CD4-CD8-thymocytes. **(D)** Graph of the percentage of CD25 and CD44 expressed CD4-CD8-cells. **(E)** Gated lin-CD25-CD44+ thymocytes stained for CD24 and CD117 expression. **(F)** Graph of the percentage of CD117+CD24high and CD117+CD24low cells in +/Z, Tg+;+/Z, Z/Z and Tg+;Z/Z mice. **(G)** FOXP3 in gated CD45+ thymocytes. **(H)** The percentage of FOXP3+ cells. **(I)** Immunostaining for FOXP3 on paraffin sections of thymus. Scale bar=100μm. n≥8

We have previously reported that within the DN population, *Z/Z* mice have increased frequency of CD44+CD25-DN1 cells relative to wild-type and heterozygotes (Fig. 4C, D) (Chen et al., 2009), due to an increase in a specific CD44+CD25lo subset (Fig. 4C, red box). This subset likely represents a partial block in DN1-DN2 differentiation, as CD44+CD25+ (DN2) and CD44-CD25+ (DN3) cells were decreased (Fig. 4C, D) (Chen et al., 2009; Xiao et al., 2018). Notably, this aberrant DN1 population was absent in *K5.Foxn1;Z/Z* mice (Fig. 4C). Both the DN1 and DN3 frequencies improved in the *K5.Foxn1;Z/Z* mice, although did not reach control levels, and DN2 frequencies did not change (Fig. 4C, D; Supplemental Fig. S4G, H). Since we did not include lineage markers, the DN1 subset in this analysis includes both c-kit+ (ETP) T lineage progenitors as well as B and NK lineage cells, all of which can be separated by their expression of HSA (CD24) and ckit (Porritt et al., 2004). Analysis of these subsets revealed that the T lineage specific CD24+ckit+ DN1a,b/ETP cell population that is decreased in *Z/Z* mutants was partially rescued in *K5.Foxn1;Z/Z* mice at 1 month (Supplementary Fig. 4I, J), and was similar to +/Z controls by 6 months (Fig. 4E, F).

We also assessed the generation of Foxp3+ regulatory T cells (T_reg_) at 6 months. T_reg_ frequencies were similar in *+/Z, Z/Z,* and *K5.Foxn1;Z/Z* thymi (Fig. 4G,H,I). Surprisingly, *K5.Foxn1;+/Z* thymi exhibited markedly increased frequencies and absolute numbers of Foxp3+ regulatory T cells (Fig. 4G, H, I, Supplemental Fig. 4D). These Treg cells expressed normal levels of CD4, TCRβ, and Foxp3 (Supplemental Fig. 5A-E). Increases in FOXP3+ cells were also seen in *K5.Foxn1* transgenics on a WT background, as FOXP3+ cells declined between 1 and 6 months in WT, but were maintained in *K5.Foxn1* transgenics at 6 months (Supplemental Fig. 5F, G). This result suggests that higher FOXN1 levels create a microenvironment that biases T cell development toward Treg generation, and that the better maintenance of mTEC phenotypes in trangenics sustained Treg generation with aging.

Taken together, these data indicate that presence of the transgene did not affect the major categories of normal thymocyte development in WT or +/*Z* mice, with the exception of increased T_reg_ production. In addition, the transgene improved thymocyte differentiation defects in *Z/Z* mice, especially increasing ETPs to levels similar to controls. This result is consistent with improved, but not normal, TEC phenotypes that better support thymocyte development.

### The *K5.Foxn1* transgene affects the differentiation and proliferation of putative TEPC

Several reports have identified Plet-1 as a marker for a population of cells with both cortical and medullary progenitor activity, both at fetal and postnatal stages (Bennett et al., 2002; Gray et al., 2006; Ulyanchenko et al., 2016). In addition, Plet1-Cld3,4^hi^ UEA1+ TECs represent progenitors for postnatal mTECs, including AIRE+ mTECs (Hamazaki et al., 2007). Given the improved maintenance of mTEC populations in the presence of the *K5.Foxn1* transgene, we evaluated the impact of the *K5.Foxn1* transgene on these progenitor populations in both wild-type mice and in the context of the *Foxn1^Z^* allele by IHC. The frequency of Plet1+ cells was significantly increased in *K5.Foxn1* transgenic mice compared to WT controls (Fig. 5A, B, Supplemental Fig. 6A, B). When all three markers were evaluated on the +/Z and Z/Z mice with and without the *K5.Foxn1* transgene, Plet1^-^Cld3^hi^UEA-1^hi^ mTEC-committed progenitors were present in all genotypes (Fig. 5C, pink arrows). In the +/*Z* thymus, most Plet1+ cells had a range of Cld3 expression levels and were UEA1- (white arrows), while a minority of these cells were also UEA-1lo (yellow arrow). Similar results were seen in the *Z/Z* mutants. In contrast, in both the *K5.Foxn1;+/Z* and *K5.Foxn1;Z/Z* thymi nearly all Plet1+ cells are both Cld3+ and UEA1+ (yellow arrows).

**Fig. 5.**
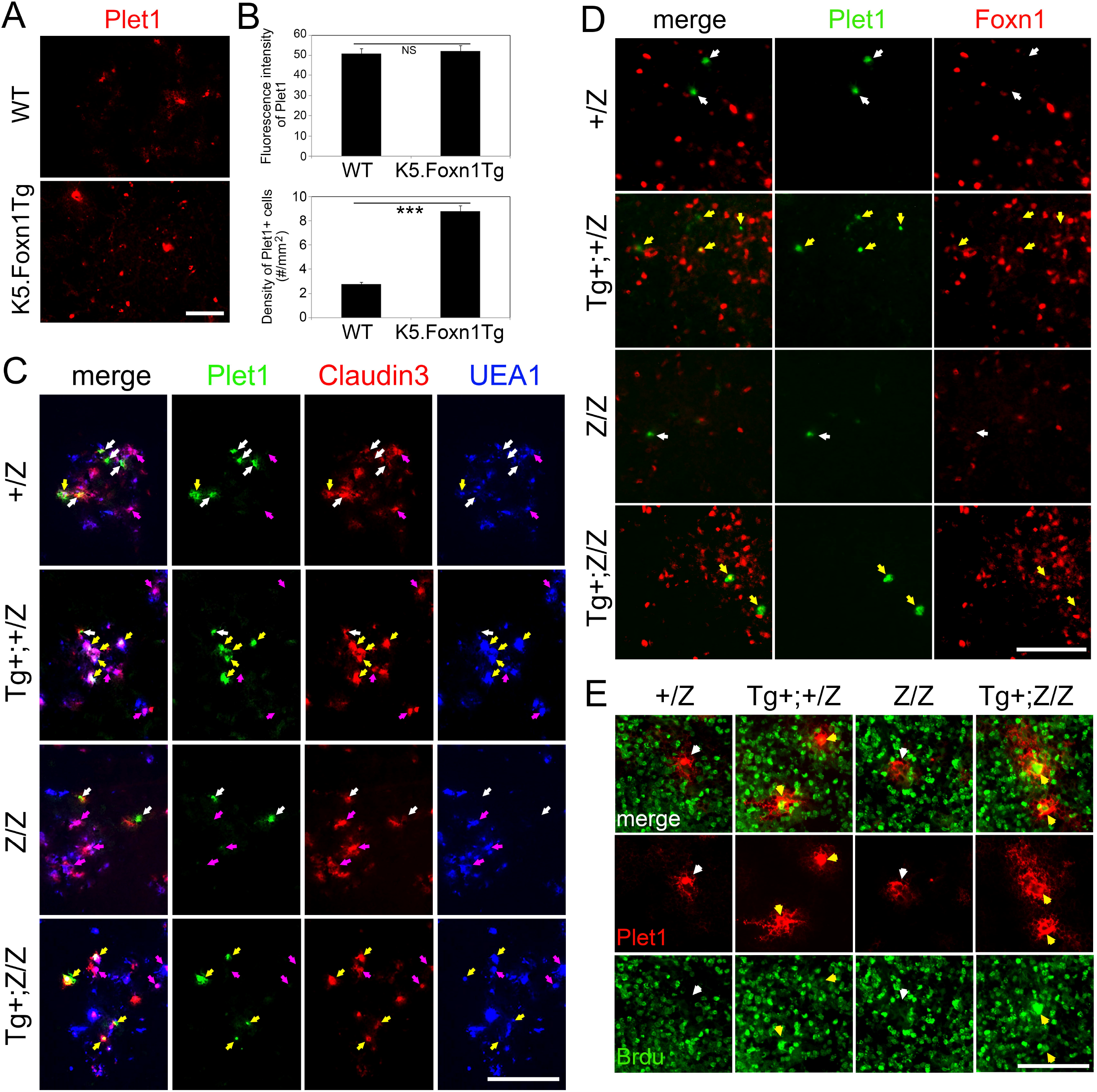
The *K5.Foxn1* transgene modulates the phenotype, frequency, and proliferation of TEC progenitors. All data from one month old mice. **(A)** PLET1+ cells (red) are increased in *K5.Foxn1* Tg thymus. **(B)** Fluorescence intensity and density of PLET1+ cells. **(C)** IHC for PLET1 (green), Claudin3 (red) and UEA-1 (blue). PLET1+Claudin3+UEA-1-cells, white arrows; PLET1+Claudin3+UEA-1+ cells, yellow arrows; PLET1-Claudin3+UEA1+ cells, pink arrows. **(D)** IHC for PLET1 (green) and FOXN1 (red). PLET1+FOXN1-cells, white arrows; PLET1+FOXN1+ cells, yellow arrowss. **(E)** IHC for PLET1 (red) and BrdU (green). PLET1+BrdU-cells, white arrows; PLET1+BrdU+ cells, yellow arrows. Scale bar=50μm. n≥3

To test whether this co-expression in the presence of the transgene correlated with presence of FOXN1, we performed IHC for FOXN1 and Plet-1. In +/*Z* and *Z/Z* mice, 90% of the Plet1^hi^ cells were negative for FOXN1 (Fig. 5D, white arrows; Supplemental Fig. 6C). In contrast, in both *K5.Foxn1;Z/+* and *K4.Foxn1;Z/Z* mice, ~50% of Plet1+ cells were Foxn1+ (Fig. 5D, yellow arrows; Supplemental Fig. 6C), although with a range of levels, indicating that the transgene was driving ectopic *Foxn1* expression in this population that contains TEC progenitors. Since *Foxn1* has been shown to be involved in the regulation of TEC proliferation (Chen et al., 2009; Itoi et al., 2007; O’Neill et al., 2016), we used BrdU incorporation to test whether increased *Foxn1* expression in these cells affected their proliferative status. We found that only 20% of Plet1+ cells in +/*Z* and *Z/Z* thymi incorporated BrdU during a 5 day period of exposure, while the majority of Plet1+ TEC in *K5.Foxn1;+/Z* and *K5.Foxn1;Z/Z* thymi were proliferating during this time window (Fig. 5E; Supplementary Fig. 6D). Additional analysis showed that there was not an overall change in TEC proliferation in either MHC^hi^ or MHC^lo^ TECs with the presence of *Foxn1* transgene, including in *Z/Z* mice that have reduced MHCII^lo^ proliferation (Chen et al., 2009) (Supplemental Fig. 6E, F). Thus, the increase in Plet1+ TEC proliferation was specific to that population.

### The *K5.Foxn1* transgene is sufficient to induce thymus development in nude mice

The *Foxn1* null thymus phenotype (nude) results from an early block in thymus development, with the resulting rudiment composed of both Plet-1+ TEPC and Cld3+UEA-1+ medullary progenitors (Depreter et al., 2008; Hamazaki et al., 2007; Nehls et al., 1996; Nowell et al., 2011). To test whether the *K5.Foxn1* transgene can rescue this phenotype, we crossed it into the nude background. In *K5.Foxn1;nu/nu* mice, the transgene was the only source of *Foxn1* since endogenous *Foxn1* is eliminated by the nude mutation (Nehls et al., 1996). Addition of the transgene to *+/nu* heterozygotes did not significantly impact thymus weight (P=0.073), the number of FOXN1+ cells, or overall cortico-medullary organization (Fig. 6A, B, E-G). IHC for K8 (cTECs), K5, and UEA-1 (mTECs) showed minor disruption of the cortical and medullary organization and regions of K8lo TECs in *K5.Foxn1Tg;+/nu* mice (Fig. 6C). Analysis of UEA-1 staining showed that addition of the transgene to *+/nu* increased frequency of UEA1+ mTECs but did not impact Cld3 staining (Fig. 6D), consistent with the results in wt and +/*Z* mice (Figs. 3, 5).

**Fig. 6.**
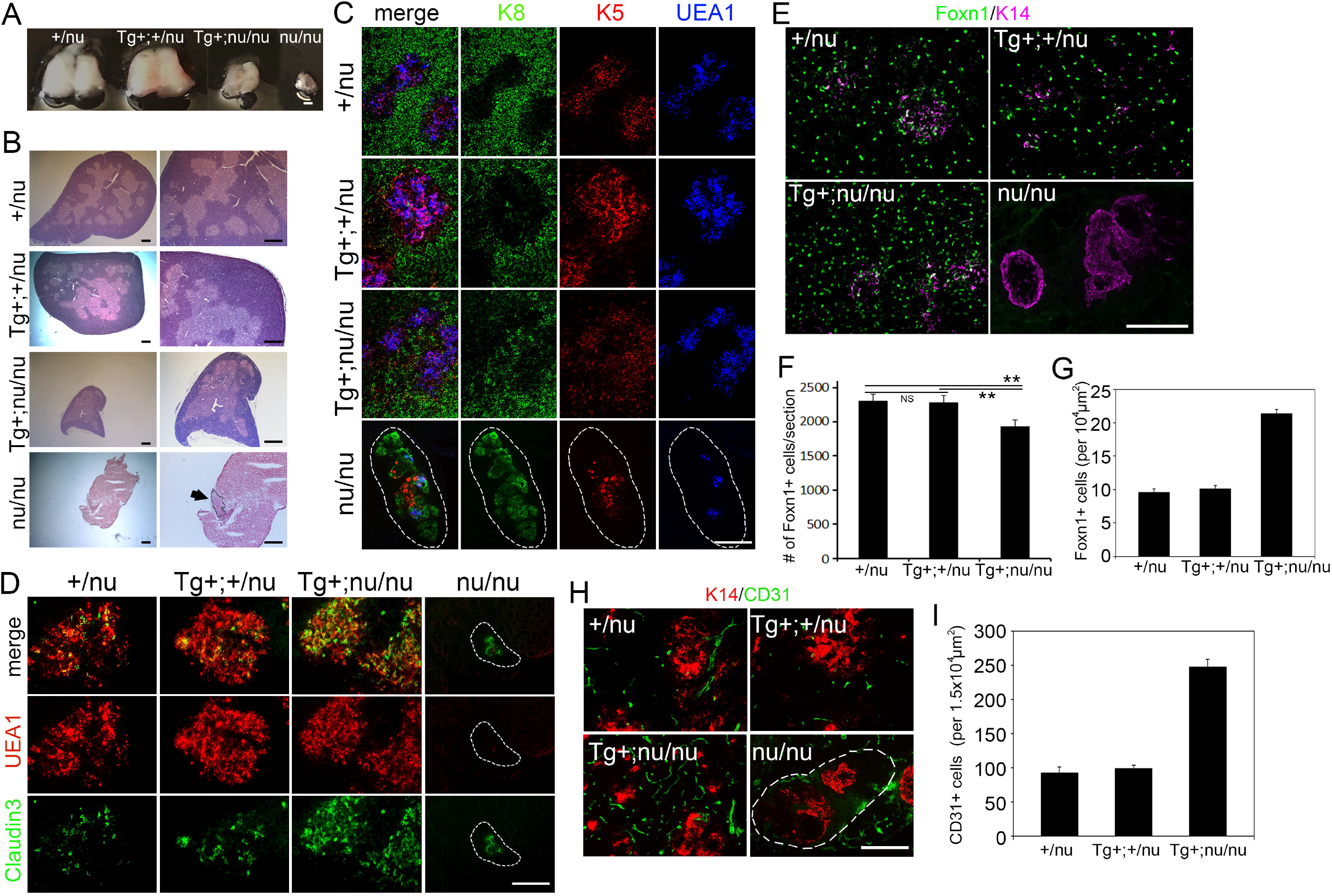
The *K5.Foxn1* transgene induces thymus development in nude mice. All data are from two week old mice. **(A)** Whole dissected thymi from mice with combinations of *Foxn1^nu^* and *K5.Foxn1Tg.* **(B)** H&E of thymi from the indicated genotypes. The right panels are a higher magnification of the left panels; arrow in the *nu/nu* right panel indicates the outlined thymic rudiment. **(C)** IHC for K8 (green), K5 (red), and UEA-1 (blue). White dotted outlines in C, D, and H indicates the thymus boundary in *nu/nu* samples. **(D)** IHC for Claudin3 (green) and UEA-1 (red). **(E)** IHC for FOXN1 (green) and K14 (pink). **(F)** Total numbers of FOXN1+ cells per section, based on cell counts of 3 sections each from the central part of 2 thymi per genotype by IHC. **(G)** Numbers of FOXN1+ cells per 10^4^μm^2^. **(H)** IHC for CD31 (green) and K14 (red). **(G)** Numbers of CD31+ cells per 1.5×10^4^μm^2^. Scale bar=100μm.

Addition of the transgene in the *nu/nu* thymus caused a dramatic improvement in thymus development. *K5.Foxn1;nu/nu* thymi were clearly larger than in *nu/nu* (Fig. 6A, B), and the number of Foxn1+ cells increased to nearly that of *+/nu* and *K5.Foxn1Tg;+/nu* mice, (Fig. 6E, F), although the density of FOXN1+ cells was increased 2-fold (Fig. 6E, G). FOXN1 expression in TECs is required to recruit endothelial cells and to generate the cellular and molecular environment needed for normal thymic vascularization (Bryson et al., 2013; Mori et al., 2010). In the nude thymus anlagen, no CD31+ cells are detected in the epithelial region (Fig. 6H). All other genotypes showed the presence of vasculature, indicating that transgene-driven TEC differentiation is capable of recruiting vasculature into the thymic rudiment. CD31+ blood vessels also had a higher density in *K5.Foxn1Tg;nu/nu* thymus compared to controls (Fig. 6H, I), consistent with the increased density of FOXN1+ TECs.

The *K5.Foxn1;nu/nu* thymus exhibited restoration of cortical and medullary compartments, with the medulla relatively expanded (Fig. 6B). Consistent with previous reports (Chen et al., 2009), *nu/nu* thymi showed broad expression of K8 with central small numbers of K5+ or UEA-1^lo^Cld3+ cells (Fig. 6C, D). *K5.Foxn1Tg;nu/nu* thymus displayed K8 expression that was similar to *K5.Foxn1Tg;+/nu* thymus, with K8^lo^ regions in the cortex and scattered K8+ cells in the medullary regions (Fig. 6C). K5 expression was throughout the medulla but also in scattered cells in the cortex that were primarily K8+ (Fig. 6C) consistent with expansion of a progenitor-containing K8+K5+ population (Klug et al., 2002). In contrast to K5 expression, UEA-1+ mTECs were localized to clearly delineated medullary areas of the *K5.Foxn1Tg;nu/nu* thymus.

Cld3+ cells are present in the nude mouse thymic rudiment, consistent with a previous report that some mTEC lineage divergence occurs in the absence of Foxn1 (Nowell et al., 2011). Our analysis confirmed that Cld3+ cells are present in the *nu/nu* thymus and are mostly UEA-1 negative or low, similar to +/nu controls (Fig. 6D). The number and frequency of Cld3+ cells was dramatically expanded in *K5.Foxn1Tg;nu/nu* mice, compared to all other genotypes, the majority of which were UEA1+. This analysis suggested that the majority of mTEC in the *K5.Foxn1Tg;nu/nu* thymus have phenotypes consistent with an mTEC progenitor (Fig. 6D).

In summary, *Foxn1* expression from the transgene is capable of driving substantial thymus differentiation and growth from the nude thymus anlage. The resulting TECs have expanded compartments of cells previously shown to contain TEC progenitors (K8+K5+ and Cld3+ cells), consistent with the transgene driving both proliferation and differentiation of these progenitors.

## Discussion

Given previous data showing that *Foxn1* overexpression in TECs attenuated and delayed thymic involution (Bredenkamp et al., 2014a; Zook et al., 2011), the fact that *Foxn1* overexpression from this *K5.Foxn1* transgene neither causes thymic overgrowth nor impacts the timing or degree of thymus size reduction during involution is surprising. However, this result is consistent with our data demonstrating that the thymus is highly sensitive to *Foxn1* dosage, as reductions in *Foxn1* mRNA levels of as little as 15% has measurable corresponding reductions in thymus size and TEC phenotypes (Chen et al., 2009). *Foxn1* over expression from the *K5.Foxn1* transgene does have specific impacts on TEC and overall thymus phenotypes with improved mTEC maintenance and expanded TEC progenitor populations on a wild-type background. In addition, this transgene improves proliferation, differentiation, and organization of both cTECs and mTECs in *Foxn11acZ* homozygous mice, including upregulation of MHCII expression in all TECs. These changes particularly impact mTEC numbers, differentiation, and maintenance. Furthermore, when expressed in *Foxn1* null nude mice the transgene drives sufficient *Foxn1* expression to promote substantial thymus development biased toward mTECs.

So why would expression of the K5.Foxn1 transgene result in a different phenotype compared to the other published accounts of *Foxn1* over-expression in TEC? There are two main differences between this transgene and the previously published accounts that could underlie this seemingly paradoxical result. First, this transgene drives *Foxn1* expression at a level that is lower than that reported for the other two transgenes. As *Foxn1* is quite dosage sensitive in loss of function models, it is certainly possible that very high levels of expression are required to cause the thymic overgrowth. Second, the promoter used in this transgene drives *Foxn1* overexpression in progenitors that are usually *Foxn1* low or negative. The two prior reports used either a K14 promoter (Zook et al., 2011) or *Foxn1^Cre^* (Bredenkamp et al., 2014a) to drive *Foxn1* expression, neither of which would drive expression in the earliest TEC progenitors. K5, in contrast, is a known marker of TEC that contain progenitor activity, as well as most or all mTECs (Klug et al., 2002; Ulyanchenko et al., 2016), and our data clearly show that this transgene drives *Foxn1* expression in Plet1+ cells that both have been implicated as progenitors, and are normally *Foxn1* negative (or below the level of detection), and results in their proliferation and relative expansion. Both features of this transgene could contribute to observed phenotypes; the expression in both bipotent Plet1+ progenitors and Cld3+ mTEC progenitors (both of which are present in the absence of *Foxn1)* could bias them towards proliferation, while the modest overexpression in immature TEC (likely contained within the MHCII^lo^ compartment) is insufficient to cause the expansion phenotypes seen under conditions of significantly higher *Foxn1* overexpression driven by the K14 or *Foxn1* promoters in other studies. This combination of effects in different TEC populations could result in the failure to observe thymus overgrowth while driving the expansion of progenitors, both in the K5.Foxn1 transgenics alone, and in combination with the *Foxn1^Z/Z^* mutation.

Other than its effects on progenitors, increased FOXN1 levels via the *K5.Foxn1* transgene had the most obvious effects on the mTEC sublineage in the postnatal thymus. In addition to causing expansion of Cldn3+ mTEC progenitors, the transgene appears to bias TEC differentiation directly toward the mTEC lineage, with increased expression of mTEC markers in the Plet1+ compartment. In addition, both UEA1 and Aire mTEC differentiation markers are up-regulated more broadly in mTECs. The up-regulation of Aire, both in numbers and on a per cell basis, is particularly important, as it could indicate an impact on self-tolerance. The efficiency of negative selection is dependent on the presentation of peripheral tissue-specific self-antigens (TSA) (Klein et al., 2014), the expression of which is in part regulated by the *Aire* gene in mTECs (Peterson et al., 2008). Postnatal reduction of *Foxn1* levels in the Z/Z mice induces loss of mTECs and reduces both the numbers of Aire+ mTECs and its expression level, both of which are increased with *Foxn1* overexpression. Although there is no evidence that *Aire* is itself a direct target of FOXN1 (Zuklys et al., 2016), these data support the idea that *Aire* expression is dependent on, and correlated with, FOXN1 levels.

An effect on negative selection is also indicated by the dramatic increase in the production of FOXP3+ Treg, which are an important component or peripheral tolerance, and are generated in the thymus by diverting cells from apoptosis during the process of negative selection. Other than this phenotype, the main effects of the transgene on thymocyte differentiation were relatively mild, although consistent with known roles for FOXN1-dependent processes. No significant differences in the main thymocyte stages defined by CD4 and CD8 were seen between +/*Z* controls and *K5.Foxn1;+/Z* trangenics; this is consistent with the relatively minor effects on TEC differentiation and the lack of any change in thymus size. However, the increased *Foxn1* expression from the transgene did fully or partially rescue the thymocyte differentiation phenotypes seen in the *Z/Z* mice. In particular, the frequency of DN1a,b/ETP progenitors was recovered consistent with its dependence on the expression of *Dl4,* a known *Foxn1* direct target, on cTECs (Zuklys et al., 2016). These results are consistent with the idea that most aspects of thymocyte differentiation are sensitive to reduced *Foxn1* dose but may be relatively insensitive to increased *Foxn1* levels.

Efforts to generate TECs via either directed differentiation or reprogramming continue to be of interest, as do targeting the *in vivo* thymus for rejuvenation via a variety of approaches towards the therapeutic goal of improving immune function after damage or with aging. The results from this study highlight the critical importance of carefully controlling both target cell type and *Foxn1* dosage in these attempts. While high levels of overexpression cause thymic overgrowth that does increase output but can be detrimental, the lower levels of expression and targeting to progenitors shown here did not result in overgrowth, despite the expansion of progenitor populations. This overexpression did maintain and improves thymic phenotypes, especially in the medulla, without overgrowth. These results demonstrate that the effects of Foxn1 overexpression are pleiotropic and differ depending on the cell type and level of expression.

## Materials and methods

### Mice

The *Foxn1^lacZ^* allele was generated by our lab (Chen et al., 2009). Nude mice (*Foxn1^nu^*) were purchased from The Jackson Laboratory Animal Resource Unit, Bar Harbor, ME. Genotyping for both alleles was as described (Chen et al., 2009). The *K5.Foxn1* transgenic line was provided by Dr. Janice L. Brissette, and was genotyped by PCR as previously reported (Weiner et al., 2007).

### Antibodies

The following antibodies were used in this study: anti–CD4 (GK1.5, Biolegend), anti-CD8 (53-6.7, Biolegend), anti-CD25 (PC61, Biolegend), anti-CD44 (IM7, Biolegend), anti-K5 (AF138, Covance), anti-K8 (provided by Dr. Ellen Riche, Texas), anti-Foxn1 (G20, Santa Cruz), anti-β5t (PD021, MLB), biotinylated UEA1 (Vector Labs), anti-CD205 (dp200, Abcam), anti-K14 (AF64, Covance), anti-CD45 (30-F11; Biolegend), anti-EpCAM (G8.8, Biolegend), anti–I-A/I-E (M5/114.15.2, Biolegend), anti-Aire (M-300, Santa Cruz), anti-CD24 (M1/69, Biolegend), anti-CD117 (2B8, Biolegend), anti-CD19 (6D5, Biolegend), anti-Foxp3 (FJK-16s, eBioscience), Plet1 (made by our Lab), anti-Claudin3 (Invitrogen), anti-BrdU (Pharmingen), anti-CD31 (PECAM-1, Pharmingen). Fluorochrome-conjugated anti-Ig second step reagents were purchased from Jackson ImmunoResearch (West Grove, PA). Binding of biotinylated Abs was detected by fluorochrome-conjugated streptavidin (Invitrogen).

### Thymic stromal cell isolation by enzymatic digestion

Thymic stromal cells were isolated as described previously (Chen et al., 2009). Briefly, adult thymi were minced, digested in collagenase (Roche, Basel, Switzerland), then collagenase/dispase (Roche) and passed through 100-μm mesh to remove debris.

### Immunohistology

Serial frozen sections (10 μm) from mouse thymus were air dried for 30 min before acetone fixation. Thin sections were blocked with normal serum and subsequently incubated with optimal dilutions of primary Abs for at least 1 hour at room temperature before washing and incubation with appropriate fluorochrome-conjugated secondary reagents. Controls included slides incubated with nonimmune species matched Ig or isotype-matched mouse Ig. For multiple Ab staining, the sections were incubated simultaneously with primary Abs from different species. Microscopic analysis was performed with a Zeiss Axioplan2 microscope or Zeiss LSM 710 (Zeiss, Melville, NY).

### Flow cytometry

Cells in PBS containing 2% FBS and 0.1% sodium azide were incubated with directly conjugated or biotinylated Abs on ice for 30 min followed by two washes. Binding of biotinylated Ab was detected with PerCp-SA. Cells were analyzed with a Cyan flow cytometer (Miami, FL) equipped with an argon laser (488 nm) for FITC and PE excitation and a heliumneon laser (633 nm) for APC and PerCp-SA excitation. Data were collected on using a four-decade log amplifier and were stored in list mode for subsequent analysis using Flowjo Software.

### Bromodeoxyuridine (BrdU) treatment and staining

Incorporation was initiated by intraperitoneal injection of BrdU (1 mg in PBS; Sigma, St Louis, MO) and maintained in drinking water for 5 days (0.8 mg/mL BrdU). Thymic stromal cells were isolated and surface labeled, then fixed in 1% paraformaldehyde, 0.01% Tween-20 (BDH Laboratory Supplies) in PBS overnight at 4°C. Cells were washed in PBS, recovered by centrifugation, and incubated in DNAse I (50 Kunitz; Roche) for 30 minutes at 37°C. After washing, cells were stained with FITC-conjugated anti-BrdU (BD Biosciences) for 1 hour at room temperature. For immune-fluorescent staining, to detect BrdU incorporation, thymic sections were incubated in 2 N HCl for 20 min at room temperature. After washing in PBS, the sections were incubated with mouse anti-BrdU (B-D Sciences, San Jose, CA) for 1 h at room temperature followed by incubation with FITC anti-mouse-IgG (Jackson ImmunoResearch).

### Quantitative PCR

Total RNA from sorted CD45-EpCam+UEA1+ and CD45-EpCam+UEA1-thymic stromal cells from 4-week-old mice were isolated using the RNEasy kit (Qiagen). cDNA was synthesized from the total RNA and used as a template for real-time relative quantitation for Foxn1 using commercially available probes and reagents (Invitrogen) on an Applied Biosystems 7500 Real Time PCR System.

### Fluorescent analysis by Image J

All fluorescent intensity and cell density on IHC were analyzed by using free software Image J (Supplemental Fig. 7)

### Statistics

Data are presented as the mean and SEM. Comparisons between two groups were made using Student’s t Test, or ANOVA for multiple group comparisons. P < 0.05 was considered significant.

---

**Supplemental Fig. 1 Thymocytes in Foxn1Tg mice. (A)** Percentage of CD4 and CD8 SP cells in one month WT and *K5.Foxn1*Tg thymocytes. **(B)** Percentage of DN1-4 subsets of CD4-CD8-cells defined by CD25 and CD44 in one month WT and *K5.Foxn1* Tg thymocytes. **(C)** Profile of CD4, CD8, CD25, and CD44 expression in *K5.Foxn1* Tg+ and WT thymocytes at six months old. **(D)** Percentage of CD4 and CD8 SP cells in six month WT and *K5.Foxn1* Tg thymocytes. **(E)** Percentage of DN1-4 subsets of CD4-CD8-cells in six month WT and *K5.Foxn1*Tg thymocytes. n≥5

**Supplemental Fig. 2 UEA1 staining in thymi from *Foxn1Z/Z* with and without the *K5.Foxn1* transgene.** All thymi are from one month old mice. **(A)** UEA-1 (green) on whole sections. The number and intensity of UEA1+ cells were decreased in the *Foxn1Z/Z* mutant. Scale bar=300μm. **(B)** Profiles of EpCam and UEA1 expression in gated CD45-thymic epithelial cells. n≥5

**Supplemental Fig. 3 The *K5.Foxn1* transgene improves Aire expression in *Foxn1Z/Z* mice.** All thymi were from one month old mice. **(A)** Total thymic epithelial cells numbers in +/Z, Tg+;+/Z, Z/Z and Tg+;Z/Z mice. **(B)** Percentage of AIRE+ cells in gated CD45-EpCam+UEA1+ thymic epithelial cells. **(C)** Gating strategy for AIRE+ cells in CD45-EpCam+UEA1+ thymic epithelial cells. n≥5

**Supplemental Fig. 4 Thymocyte phenotypes in one and 6 month old *Foxn1Z/Z* with and without the *K5.Foxn1* transgene..** Samples were from six month (A-D) and one month (E-J) mice. **(A)** Total thymocyte numbers in +/Z, Tg+;+/Z, Z/Z and Tg+;Z/Z mice. **(B)** Percentage of CD4-8- and CD4+8+ thymocytes in total thymocytes. **(C)** Numbers of CD4-8-, CD4+8+, CD4+ and CD8+ cells in thymus. **(D)** Total number of Foxp3+ cells in thymus. n≥8 **(E)** Profile of CD4 and CD8 expression in +/Z, Tg+;+/Z, Z/Z and Tg+;Z/Z thymocytes. **(F)** Percentage of CD4 and CD8 SP cells in +/Z, Tg+;+/Z, Z/Z and Tg+;Z/Z thymocytes. **(G)** Profile of CD25 and CD44 expression in gated CD4-CD8-thymocytes. **(H)** Percentages of CD4-CD8-subsets defined by CD25 and CD44 expression. **(I)** Gated lin-CD25-CD44+ thymocytes stained for CD24 and CD117 expression. **(J)** Percentages of CD117+CD24high and CD117+CD24low cells in +/Z, Tg+;+/Z, Z/Z and Tg+;Z/Z mice. n≥5

**Supplemental Fig. 5 Characterization of Foxp3+ thymocytes from *Foxn1Z/Z* thymus with and without the *K5.Foxn1* transgene. A-E** are from 6 month thymi. **(A-B)** Profiles of gated Foxp3+ cells shows similar MFI of CD4 **(A)** and TCRβ (**B**). **(C)** Profile of CD25 expression in gated Foxp3+ thymocytes. **(D)** Profile of Foxp3 expression in gated CD4+ thymocytes. **(E)** Profile of CD25 and Foxp3 expression in gated CD4+ thymocytes. **(F)** Fluorescent immunostaining offor FOXP3 on one month and six month old thymus. **(G)** Fluorescent intensity of FOXP3 analyzed by ImageJ. Scale bar=100μm. n≥3

**Supplemental Fig. 6 Characterization of PLET1 and MHCII on TECs from *Foxn1Z/Z* thymus with and without the *K5.Foxn1* transgene. (A)** Profile of PLET1 expression and **(B)** percentage of PLET1+ cells in CD45-EpCam+ WT and *K5.Foxn1* thymic epithelial cells. **(C)** Percentages of PLET1+FOXN1+ cells in PLET1+ thymic epithelial cells and **(D)** of PLET1+BrdU+ cells in PLET1+ thymic epithelial cells qualtified from IHC shown in Figure 5. **(E)** Thymic epithelial cells from one month old BrdU treated mice stained for BrdU and MHCII. **(F)** Percentage of BrdU+ populations in MHC^hi^ and MHC^lo^ thymic epithelial cells from panel E. n≥7

**Supplemental Fig. 7 The measurement of fluorescent intensity on IHC results.** Left panel is the original IHC of Foxn1, middle panel shows how ImageJ recognizes each cell with different shapes. The numbers on the right panel are the fluorescent intensity automatically generated by ImageJ.

## Notes

### Competing Interest Statement

The authors have declared no competing interest.

